# Overburdened Ferroptotic Stress Impairs Tooth Morphogenesis

**DOI:** 10.1101/2023.06.08.544273

**Authors:** H.S. Wang, X.F. Wang, L.Y. Huang, C.L. Wang, F.Y. Yu, L. Ye

## Abstract

The role of regulated cell death (RCD) in organ development, particularly the impact of non-apoptotic cell death, remains largely uncharted. Ferroptosis, a non-apoptotic cell death pathway known for its iron dependence and lethal lipid peroxidation, is currently being rigorously investigated for its pathological functions. The balance between ferroptotic stress (iron and iron-dependent lipid peroxidation) and ferroptosis supervising pathways (anti-lipid peroxidation systems) serves as the key mechanism regulating the activation of ferroptosis. Comparing to other forms of regulated necrotic cell death (RNCD), ferroptosis is critically related to the metabolism of lipid and iron which are also important in organ development. In our study, we examined the role of ferroptosis in organogenesis using an *ex vivo* tooth germ culture model, investigating the presence and impact of ferroptotic stress on tooth germ development. Our findings revealed that ferroptotic stress increased during tooth development, while the expression of Gpx4, a crucial anti-lipid peroxidation enzyme, also escalated in dental epithelium/mesenchyme cells. The inhibition of ferroptosis was found to partially rescue erastin-impaired tooth morphogenesis. Our results suggest that while ferroptotic stress is present during tooth organogenesis, its effects are efficaciously controlled by the subsequent upregulation of Gpx4. Notably, an overabundance of ferroptotic stress, as induced by erastin, suppresses tooth morphogenesis.

## Introduction

A delicate balance between cell division and regulated cell death (RCD) is crucial for the development of tissues and organs [1]. In organs such as the nervous, immune, and reproductive systems, cells are initially produced in excess and are subsequently removed by RCD. During organ development, structures with transient functions are also eliminated by RCD when they are no longer necessary [2]. The best-known example of this is the formation of digits in higher vertebrates, where the interdigital webs are eliminated primarily through the apoptotic machinery [3]. RCD plays a vital role in animal development and adult life by eliminating abnormal and potentially harmful cells [4]. Apoptosis is the most extensively studied form of RCD which shrinks the nucleus and buds the plasma membrane (without rupturing it), making it essential in achieving cell number regulation, tissue remodeling, and sculpting structures, driving morphogenesis during organ development [5].

However, several new types of cell death, including pyroptosis, necroptosis, and ferroptosis, have been discovered [6]. Unlike apoptosis, these newly discovered RCDs are characterized by pore formation and/or rupture of the plasma membrane, thus termed as regulated necrotic cell death (RNCD) [7]. While apoptosis benefits organ morphogenesis in certain circumstances, whether RNCD and its regulatory mechanisms are biologically required in organ development remains poorly understood. Pyroptosis and necroptosis are mainly activated by external pathogens or damage-associated molecular patterns [8], whereas ferroptosis is a distinct form of iron-dependent, lipid peroxidation-driven programmed cell death [9]. Ferroptosis is a particular type of cell death that links lipid metabolism, ROS (reactive oxygen species) biology, iron regulation, and disease [10]. Despite the pathological role of ferroptosis and/or ferroptotic stress in multiple diseases [11,12], recent studies have identified the occurrence of ferroptosis in embryonal erythropoiesis and aging [13] and aged skeletal muscle [14], indicating its unrecognized role in physiological processes. Unlike autonomous activation of apoptosis during organ development, the broken balance between ferroptotic stress and its suppressing mechanisms is now considered the main reason for ferroptosis. Recent studies demonstrated that MYSM1 deficiency causes human hematopoietic stem cell loss by ferroptosis, highlighting the broader developmental and regenerative role of ferroptosis [15]. The investigation into the potential involvement of ferroptosis and/or ferroptotic stress, other than rest forms of non-apoptotic cell death like pyroptosis and necroptosis, in organ morphogenesis is an area of great interest. However, conducting high-throughput analysis through *in vivo* studies presents challenges as it relies heavily on generating specific transgenic mice, since erastin, the classic small molecule used to induce ferroptosis is feasible *in vivo* [16]. Thus, there’s a crucial need for a more adequate model to explore the ferroptosis/ferroptotic stress involved in development.

The process of tooth development, which contains cell number regulation (proliferation, differentiation, and extinction of ameloblast), tissue remodeling (elimination of transient structure called enamel knot), and sculpting structures (cusp formation) [17,18], is an ideal model for the investigation of RCD in organ development. Moreover, *ex vivo* culture of tooth germ is a well-established study system allows for the controlled manipulation of environmental, genetic and pharmacological factors (like erastin), which can influence developmental events of tooth [19], providing compelling evidence for its usefulness as an efficient research model in studying ferroptosis/ferroptotic stress within developmental process [20].

In the present study, erastin was used in an *ex vivo* culture model of tooth germ to investigate the possible role of ferroptosis during tooth morphogenesis. To definitively identify ferroptosis, multiple markers are needed, including lipid peroxidation, iron accumulation, mitochondria injury, and the expression of ferroptosis-related genes [9]. According to our results, Gpx4 (glutathione peroxidase 4, a peroxidase to counter the oxidation of lipids in membranes) [21] is upregulated according to odontogenesis and amelogenesis together with obvious accumulation of iron. Erastin significantly suppressed the morphogenesis of cultured tooth germ and showed a dose-dependent manner. And meanwhile, erastin-treated tooth germ has elevated lipid peroxidation indicated by expression of 4-HNE (4-hydroxy-2-nonenal, end-product of lipid peroxidation), enhanced iron accumulation, shrunken mitochondria, and alternated expression of ferroptosis related genes. However, all these phenomena could be partially rescued by Fer-1, a classical inhibitor of ferroptosis.

In summary, our results suggest that erastin induces morphologic defects of tooth germ is conducted by the activation of ferroptosis. Ferroptotic stress and its regulatory mechanism participate in tooth morphogenesis, which borders our understanding of the physiological function of non-apoptotic regulated cell death in organ development.

## Results

### Spatiotemporal characterization of Gpx4 expression and iron accumulation in tooth morphogenesis

In the different developmental stages of the mouse first mandibular molar, the expression of Gpx4 was detected by IHC (Fig 1A). These data revealed gradually upregulated expression of Gpx4 in tooth germ from E13.5 to E17.5 (Fig 1A-a to 1A-c), but started to decrease from PN1 to PN3 (Fig 1A-d to 1A-e). An enlarged view of each group indicated a different expression level of Gpx4 between dental epithelia and mesenchyme (Fig 1A-a’ to 1A-e’). After calculating the IOD (Integrated optical density) of Gpx4 in epithelia, mesenchyme, and total area of tooth germ respectively, we found that Gpx4 in dental epithelia is higher than that in mesenchyme although they both increased from E13.5 to E17.5, which could be observed in Fig 1A-a’ to 1A-d’. Negative controls are listed in Fig S1A.

**Figure 1.**
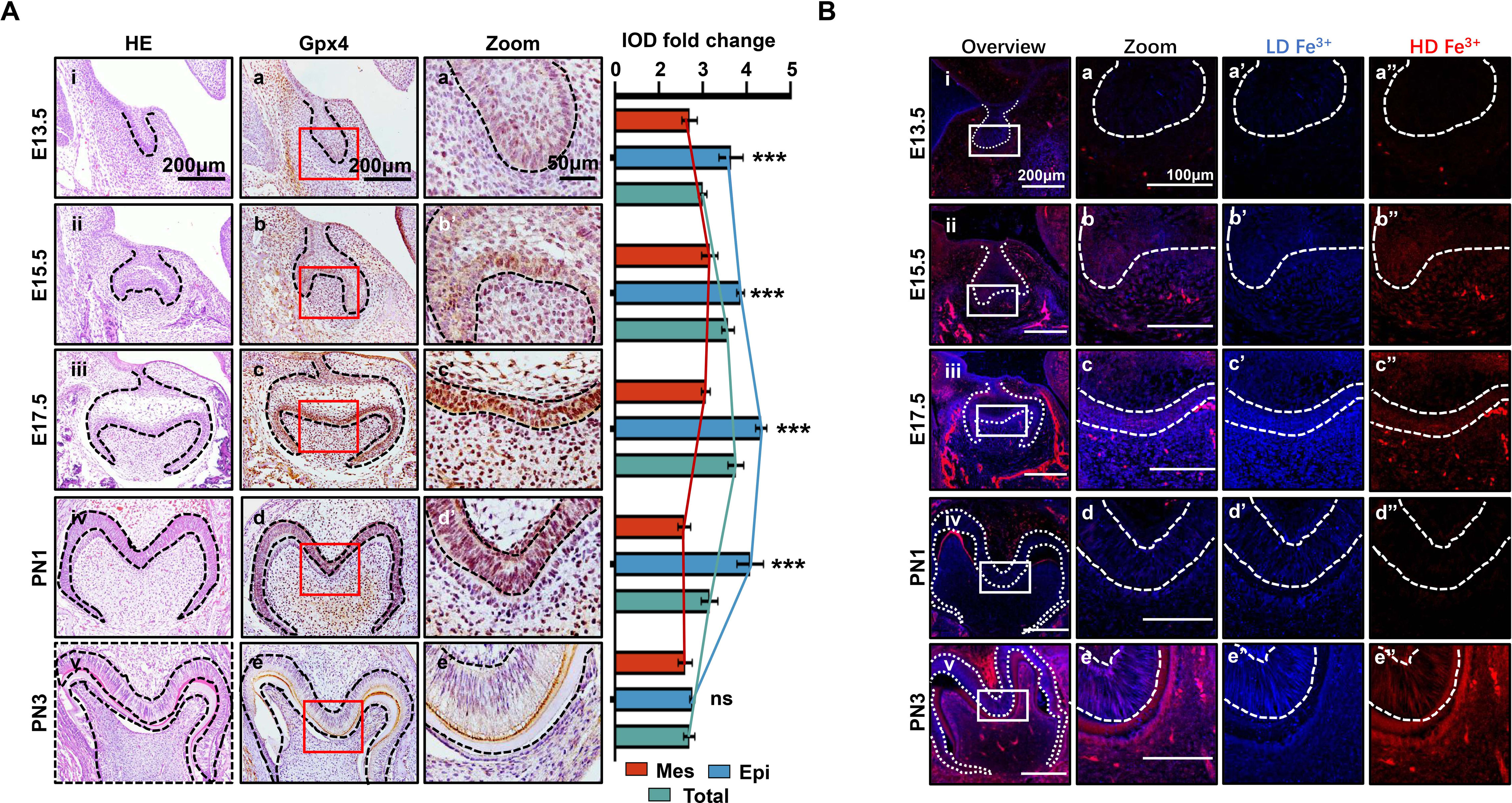
Spatiotemporal characterization of Gpx4 expression and iron accumulation in tooth morphogenesis. **(A)** (i∼v) HE staining for tooth germ in E13.5 to PN3, scar bars=200μm; (a∼e) Gpx4 expression detected by IHC, scar bars=200μm; (a’∼e’) Enlarged view of each Gpx4 staining, scar bars=50μm; epithelia versus mesenchyme *\*\*\*P<0.001* **(B)** (i∼v) Iron probe staining in for tooth germ in E13.5 to PN3, scar bars=200μm; (a∼e) Enlarged view of selected region, scar bars=100μm, low concentration (a’∼e’, blue) and high concentration of iron (a’’∼e’’, red) are present, scar bars=100μm. Epi=Epithelia, Mes=Mesenchyme.

Iron accumulation, one of the core risk factors for ferroptosis, were detected by TPE-o-Py (*ortho*-substituted pyridinyl-functionalized tetraphenylethylene) [22] using IF (Fig 1B). The results showed accumulation of iron is upregulated within dental epithelium and mesenchymal cells during differentiation of odontoblast and ameloblast (Fig 1B-a to 1B-c) then decreased at PN1 which is consistent to the expression pattern of Gpx4. However, at PN3, the active secretory period of both ameloblast and odontoblast, the accumulation of iron rebound to a high level and distributed within enamel (Fig 1B-e).

Collectively, our results demonstrated that at early stage of tooth development, enhancing risk factors of ferroptosis (iron accumulation) was accompanied by strengthening anti-ferroptosis mechanism (upregulation of Gpx4), the accumulation of iron and the expression of Gpx4 in dental epithelium/mesenchymal cells and extracellular matrix are critically related to the developmental stage.

### Erastin impairs tooth morphogenesis especially within dental mesenchyme

To investigate the possible function of ferroptosis in tooth development, an *ex vivo* culture model of the molar germ was established as described before [23], with or without treatment of erastin. Successful *ex vivo* culturing of tooth germ from D0 to D7 is presented in Fig S1B. All the molar germs were dissected from E15.5 mousses’ mandibles and cultured in medium with or without erastin (10μM) for 1, 3, and 5 days (Fig 2A). Comparing Fig 2A-d and 2A-h, gross anatomy showed apparent tiny tooth germ in the erastin treated group than that of CTRL (Control group) in D5 (Day 5). Histological analysis of D5 was performed in Fig 2B; treatment of erastin (10μM) elevated the number of necrotic-like cells (NLCs), especially in dental mesenchyme (Fig 2B-b’ and -d’). We also found a dose-dependent manner of erastin in suppressing tooth morphogenesis (Fig 2C). In D5 incubation of different concentrations (1.5, 5, 10, and 20μM) of erastin, tooth germ deteriorated morphologically (decreasing size, Fig 2C-a to -e) and histologically (elevating number of NLCs within dental mesenchyme, Fig 2C-f’ to -j’) according to the increasing concentration of erastin of erastin treatment. Relative development suppression of tooth germs was then calculated by height, width, and coronal area compared to the CTRL (Fig 2D); The radar graph showed significant dose-dependent impairment of erastin to tooth germ, the original bar graphs are listed in Fig S2.

**Figure 2.**
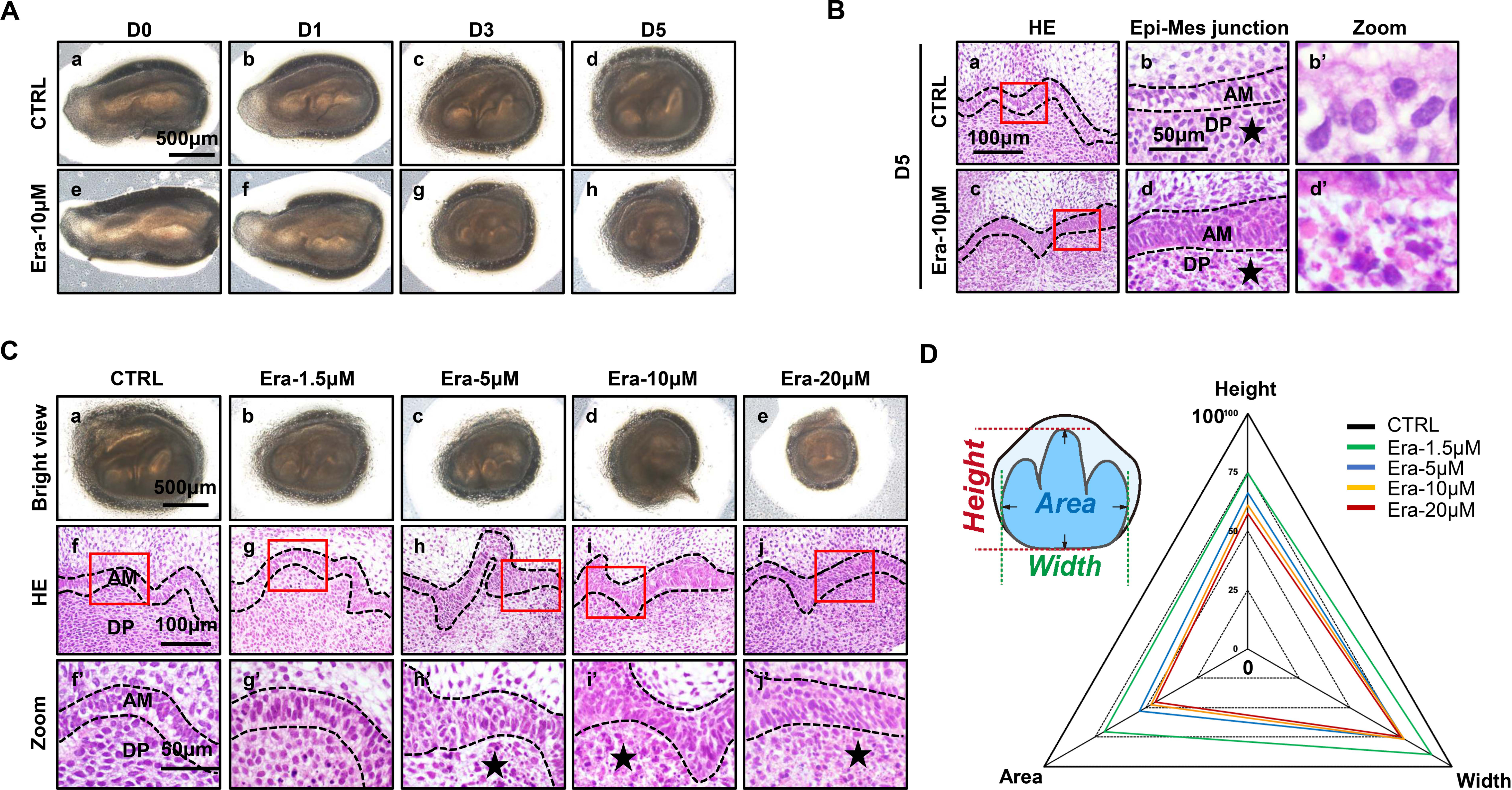
Erastin impairs tooth morphogenesis especially within dental mesenchyme. **(A)** Gross anatomy of tooth germ cultured *ex vivo* for five days, scar bars=500μm; **(B)** (a, c) HE staining for tooth germ in D5, scar bars=100μm; (b, d) High resolution of epi-mes junction papilla, scar bars=50μm; Black stars pointe out NLCs; (b’, d’) NLCs indicated by black stars. **(C)** (a∼e) Gross anatomy of tooth germs in different concentrations of erastin in D5, scar bars=500μm; (f∼j) HE staining of different concentration treated tooth germ in D5; (f’∼j’) High-resolution view of epi-mes junction region of each tooth germ, scar bars=50μm; **(D)** Rada graph for calculation of height, width, and area in each tooth germ. Black dotted line outlines ameloblasts, AM=ameloblast, DP=dental papilla.

### Ferroptosis is activated in erastin-treated molar germ

Accurate identification of ferroptosis should be determined by series markers, including iron concentration, lipid peroxidation, mitochondria dysmorphia, and overexpression of ferroptotic genes. We estimated the activation of ferroptosis in tooth germ. Indicated by TPE-o-Py detection in Era-10μM of D5 (Fig 3A-a to 3A-b’’), more iron is accumulated than that in CTRL and the region of high concentration mainly located in dental mesenchyme (red, Fig 3A-a and 3A-b). Lipid peroxidation was indicated by the expression of 4-HNE; the distribution pattern of 4-HNE was similar to iron accumulation which suggested the activation of iron-dependent lipid peroxidation occurred within Era-10μM of D5 (Fig 3A-c to 3A-d’’). Results of D1 and D3 showed similar patterns in the accumulation of iron and the expression of 4-HNE (Fig S3), also a dose-dependent manner in D5 (Fig S4). Morphological changes of mitochondria revealed severe shrinkage of mitochondria in dental mesenchyme of Era-10μM (Fig 3B). The size of mitochondria in each group is calculated in Fig 3C; results clearly showed the main size distribution of Era-10μM is significantly decreased. Then, the expression of *Gpx4*, *Slc7a11,* and *Ptgs2* in each tooth germ was modulated by q-PCR. Fig 3D showed *Ptgs2*, a gene representing the peroxidation level, dramatically increased in tooth germ after erastin treatment, while *Gpx4* and *Slc7a11*, anti-ferroptosis related genes, underwent upregulated but much more moderate expression. All these results contributed to the confirmation of the activated ferroptosis occurred in erastin-treated molar germ, and demonstrated that no lethal concentration of erastin could lead to tooth morphogenesis partially through activation of ferroptosis in dental mesenchyme.

**Figure 3.**
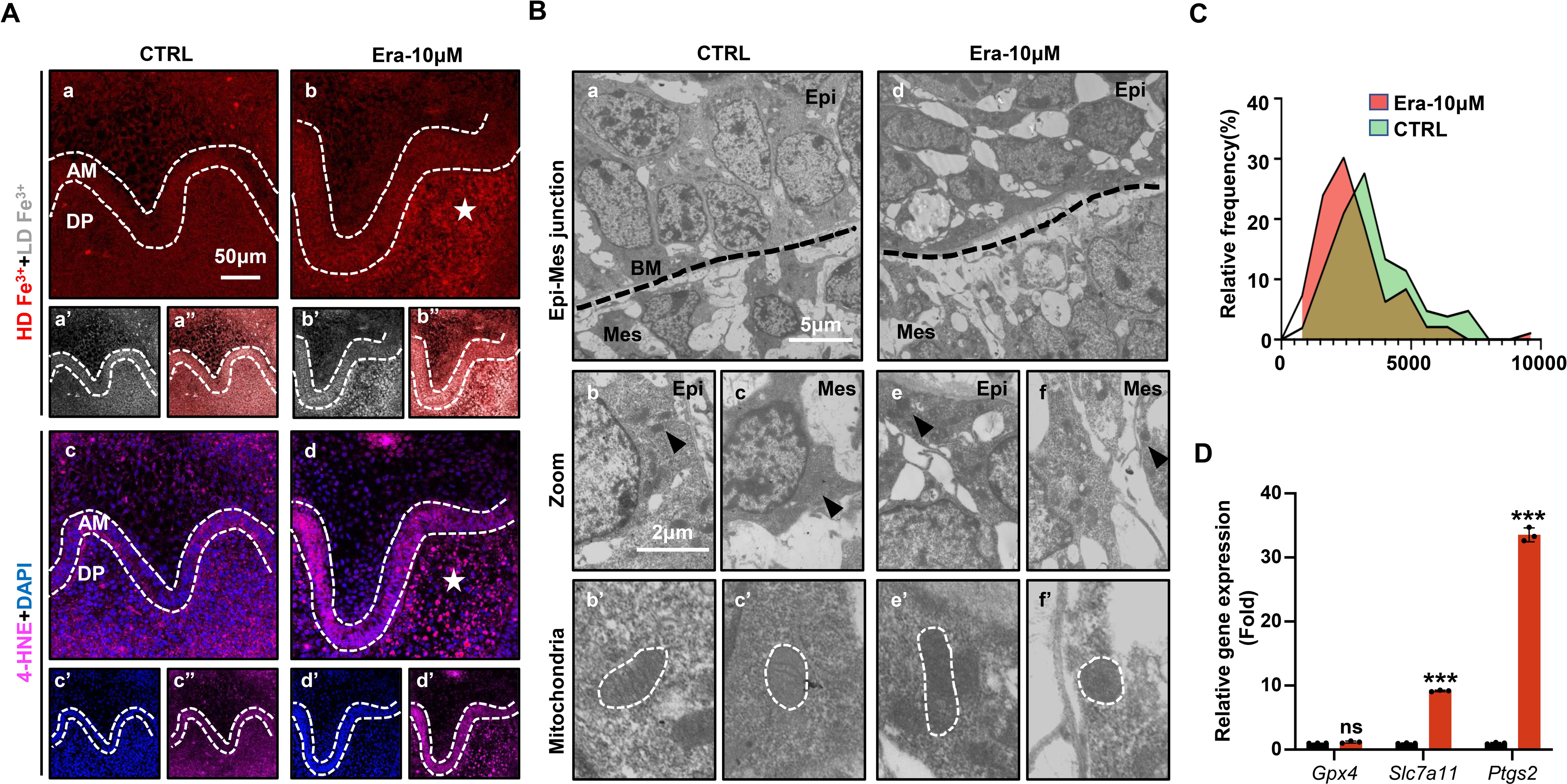
Ferroptosis is activated in erastin-treated molar germ. **(A)** (a, b) High density of Fe^3+^ (red) in CTRL and Era-10μM of D5, white star pointed out strong fluoresce signal of Fe^3+^, scar bars=50μm; (a’, b’) Low density of Fe^3+^ (grey) and (a’’, b’’) merged view of iron probe staining; (c, d) Merged view of IF straining of 4-HNE (Magenta) and DAPI (blue), white star pointed out strong fluoresce signal of 4-HNE, scar bars=5050μm; (c’, d’) for DAPI and (c’’, d’’) for 4-HNE; AM=ameloblast, DP=dental papilla, HD=high density, LD=low density; **(B)** TEM **s**canning for CTRL and Era-10μM in D5; (a, d) Epi-Mes junction area of CTRL and Era-10μM in D5 are detected, scar bars=5μm; (b, c) representative view of cells in epithelia and mesenchyme for CTRL, scar bars=2μm; Black arrow pointed out typical mitochondria in each region (b’) for epithelia and (c’) of mesenchyme, outlined by white dotted line; (e, f) representative view of cells in epithelia and mesenchyme for Era-10μM, scar bars=2μm; Black arrow pointed out typical mitochondria in each region (e’) for epithelia and (f’) of mesenchyme, outlined by white dotted line; **(C)** Relative frequency of the mitochondrial size in both groups; (D) Fold changes of gene expression in CTRL and Era-10μM in D5, versus CTRL *\*\*\*P<0.001*.

**Figure 4.**
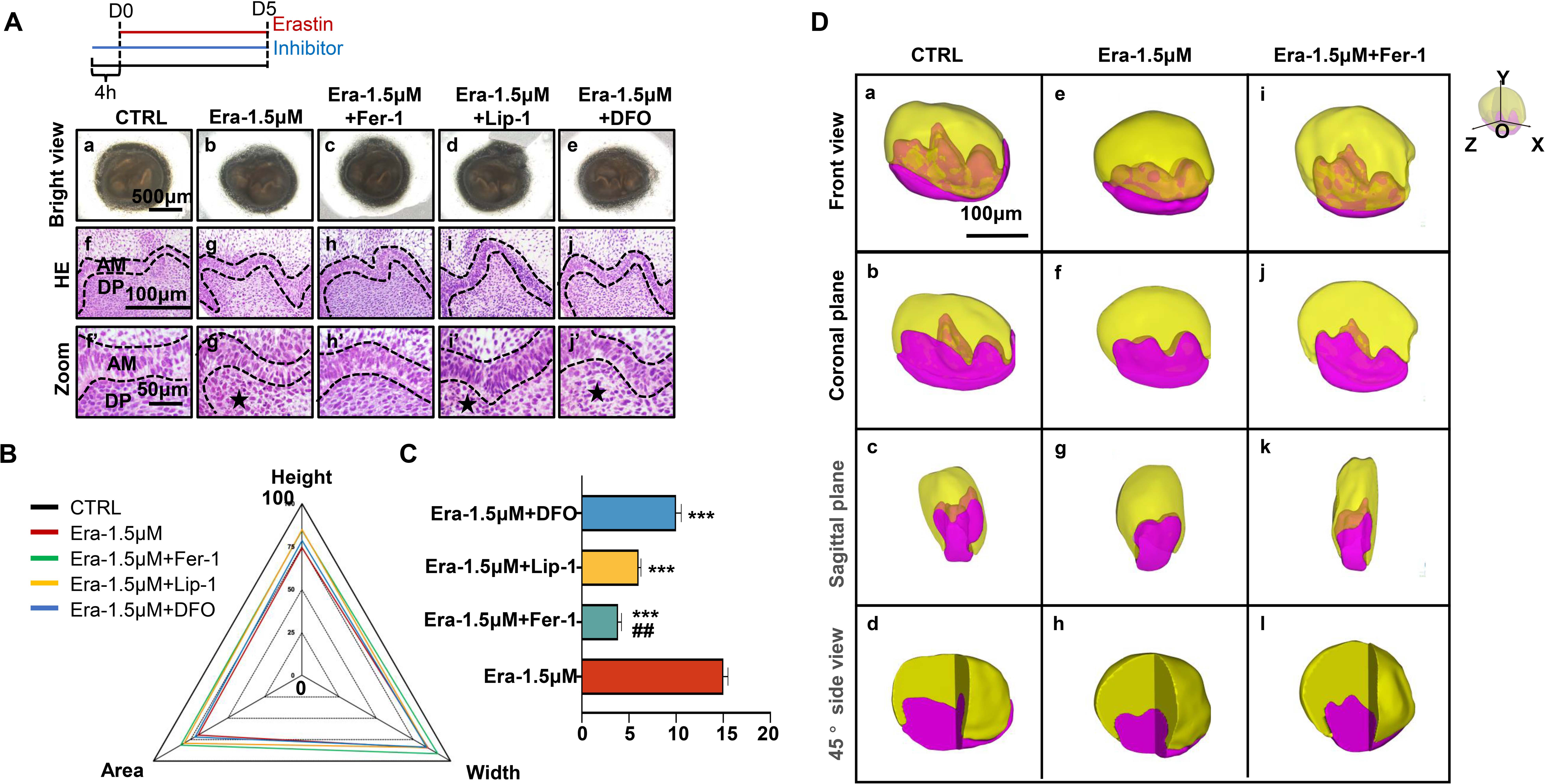
Ferroptotic inhibitors partially rescue erastin-impaired tooth organogenesis. **(A)** (a∼e) Gross anatomy of tooth germs in differently treated group, scar bars=500μm; (f∼j) HE staining of differently treated tooth germ, scar bars=100μm; (f’∼j’) High resolution view of epi-mes junction region of each tooth germ; Black dotted line outlines ameloblasts, black star pointed out NLCs, AM=ameloblast, DP=dental papilla, scar bars=50μm; **(B)** Rada graph for calculation of height, width, and area in each tooth germ; **(C)** Average number of NLCs in each group, versus Era-1.5μM *\*\*\*P<0.001*, versus Era-1.5μM+Lip-1 *##P<0.01*; **(D)** 3D reconstructed view of tooth germ in D5. (a∼d) CTRL from the front view, coronal plane, sagittal plane, and 45° side view; (e∼h) to Era-1.5μM and (i∼l) to Era-1.5μM+Fer-1 were viewed by the same way. Scar bars=100μm.

### Ferroptotic inhibitors partially rescue erastin-impaired tooth organogenesis

To further prove the function of ferroptosis in the suppression of tooth germ morphogenesis, a rescue assay was applied by using Fer-1 (classical inhibitor of ferroptosis), Lip-1 (inhibitor of lipid peroxidation), and DFO (iron chelator) to co-incubated tooth germ with erastin.

As results shown in Fig 4A, although all these three molecules show rescuing efficiency, Fer-1 holds the highest efficiency in recovering the organogenesis of molar germ in gross anatomy (Fig 4A-c) and reducing the number of NLCs at the histological level (Fig 4A-h and 4A-h’). Calculating the height, width, area of tooth germ (Fig 4B, original bar graphs are listed in Fig S5), and the number of NLCs (Fig 4C) showed inhibitors could partially rescue impairment by erastin, while Fer-1 rescued most.

**Figure 5.**
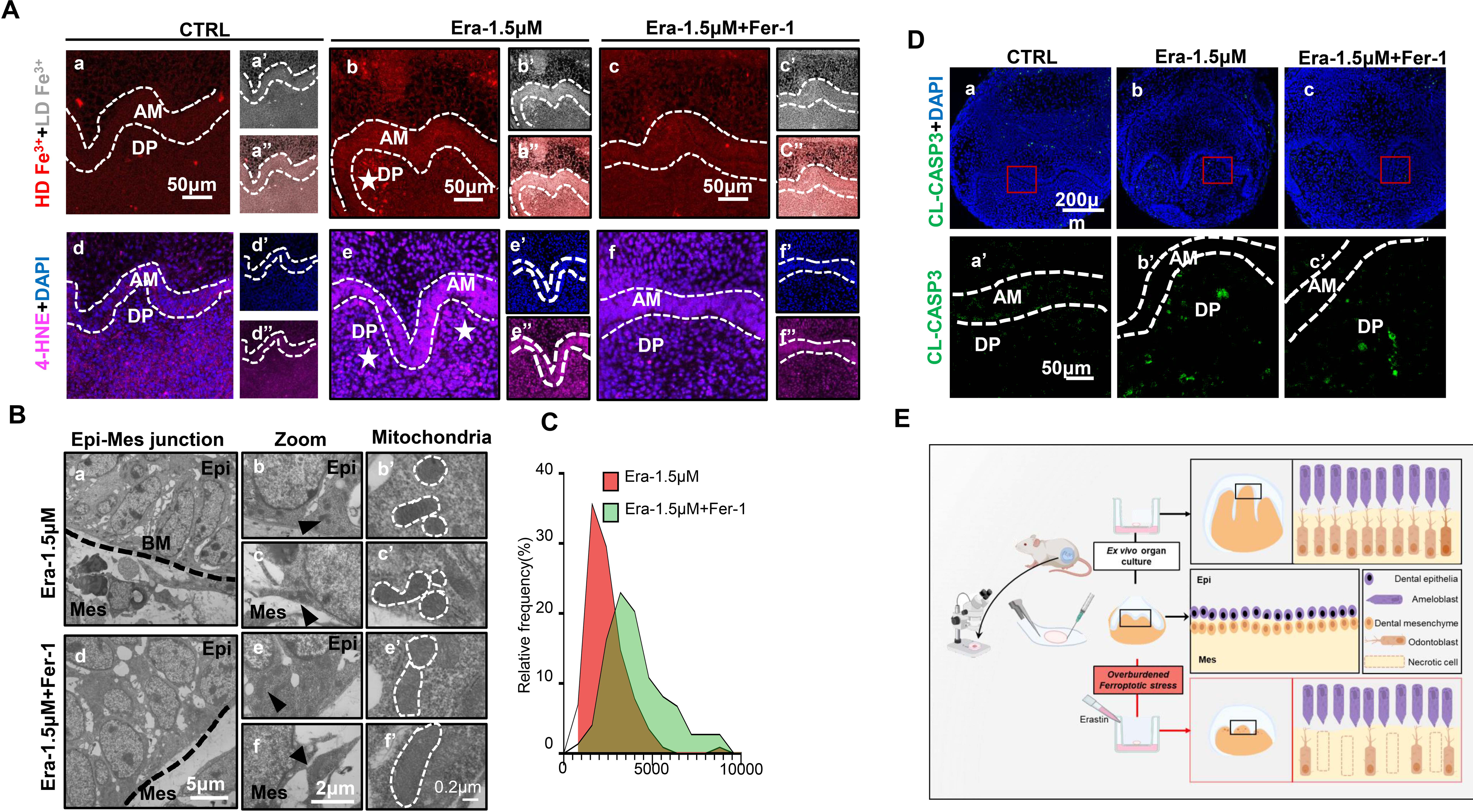
Ferroptosis is the dominant cell death type contributes to erastin-impaired tooth morphogenesis. **(A)** (a∼c) High density of Fe^3+^ (red) in CTRL, Era-1.5μM and Era-1.5μM+Fer-1 of D5, white star pointed out strong fluoresce signal of Fe^3+^, scar bars=50μm; (a’∼c’) Low density of Fe^3+^ (grey) and (a’’∼c’’) merged view of iron probe staining; (d∼f) Merged view of IF straining of 4-HNE (Magenta) and DAPI (blue), white star pointed out strong fluoresce signal of 4-HNE, scar bars=50μm; (d’∼ f’) for DAPI and (d’’∼ f’’) for 4-HNE; AM=ameloblast, DP=dental papilla, HD=high density, LD=low density; **(B)** TEM **s**canning for Era-1.5μM and Era-1.5μM+Fer-1 in D5; (a, d) Epi-Mes junction area of Era-1.5μM and Era-1.5μM+Fer-1 in D5 are detected, scar bars=5μm; (b, c) representative view of cells in epithelia and mesenchyme for Era-1.5μM, scar bars=2μm; Black arrow pointed out typical mitochondria in each region (b’) for epithelia and (c’) of mesenchyme, outlined by white dotted line; (e, f) representative view of cells in epithelia and mesenchyme for Era-1.5μM+Fer-1, scar bars=2μm; Black arrow pointed out typical mitochondria in each region (e’) for epithelia and (f’) of mesenchyme, outlined by white dotted line; **(C)** Relative frequency of the mitochondrial size in both groups. **(D)** (a∼c) Expression of CL-CASP3 (green) in CTRL, Era-1.5μM and Era-1.5μM+Fer-1 in D5, scar bars=200μm; (a’∼c’) Enlarged view of CL-CASP3 in each group, scar bars=50μm. **(E)** Schematic model illustrates the overburdened ferroptotic stress impaired tooth morphogenesis in *ex vivo* organ culture model.

To avoid bias caused by sectioning, sequential HE slides of CTRL and Era-1.5μM were applied for 3D reconstruction as previously described [24]. In Fig 5D-a to 5D-j, tooth germ in CTRL showed well-developed morphology in the 3D models. However, cusp formation in Era-1.5μM (Fig 4D-e to 4D-h) underwent significant suppression comparing to CTRL in different directions of view; original sequential slides are listed in Fig S6 to CTRL and Fig S7 to Era-1.5μM. 3D reconstruction was also performed in Era-1.5μM+Fer-1 (Fig 4D-i to 4D-l). Compared to Era-1.5μM (Fig 4D-e to 4D-h), Fer-1 reversed abnormal features dramatically in tooth morphology and cusp formation. Source sequential sections are listed in Fig S8.

### Ferroptosis is the dominant cell death type contributes to erastin-impaired tooth morphogenesis

We further estimated key characteristics of ferroptosis to further assess whether Fer-1 rescued erastin-impaired tooth germ through inhibiting ferroptosis. Comparing to CTRL group after 5 days *ex vivo* culture (Fig 5A-a and 5A-d), iron accumulation and lipid peroxidation (indicated by 4-HNE) in Era-1.5μM is much higher (Fig 5A-b and 5A-e). Fer-1 treatment vanished iron accumulation and lipid peroxidation caused by erastin (Fig 5A-c and 5A-f). Results of TEM clearly showed decreased mitochondria shrinkage in Era-1.5μM+Fer-1 than that of Era-1.5μM (Fig 5B and 5C). All these data convinced that Fer-1 rescued erastin-impaired tooth morphogenesis through inhibiting ferroptosis.

Apoptosis is the main innate cell death type in physiological process. To exclude possible over activation of apoptosis induced by erastin treatment, we detected CL-CASP3 by IF staining, main executor protein of apoptosis, in each group. In Fig 5D, CL-CASP3 is weakly expressed in CTRL group since the activation of apoptosis is physiologically required in tooth development (Fig 5D). The expression of CL-CASP3 slightly increased in both Era-1.5μM and Era-1.5μM+Fer-1, but showed no statistical differences comparing to CTRL group (Fig S9A). TUNEL assay is also performed to identify apoptotic cells by detecting DNA damage, results further convinced that apoptosis is not significantly activated in erastin treated tooth germ (Fig S9B). Taken together, ferroptosis is the dominant cell death led by erastin treatment in impaired tooth germ.

## Discussion

In the last several decades, the beneficial role of apoptosis in regulating organ development and tissue regeneration has been identified [25]. Apoptosis in tooth development had been characterized by the activation of Caspase 3 and DNA damage, which revealed a spatiotemporal apoptotic cell death pattern due to different stage of tooth morphogenesis [26]. This raises the question of whether other newly determined non-apoptotic cell death pathways also participate in the physiological processes like development, maintaining homeostasis, ageing, etc. Except for NETosis, a neutrophil extracellular trap-related cell death, the possible function and mechanism of the rest non-apoptotic cell death including, pyroptosis, necroptosis, and ferroptosis, are still barely investigated. Characterized by its close relationship with the metabolism of lipid, iron, and ROS [27,28], risk factors inducing ferroptotic stress and/or activating ferroptosis also critically involved in development and ageing, which made ferroptosis and its regulatory mechanism, other than apoptosis, pyroptosis, and necroptosis, a promising undiscovered type of cell death during tooth development.

To explore the potential involvement of ferroptosis in organogenesis, we have developed an *ex vivo* culture model of tooth germ morphogenesis. This system allows for the application of erastin to induce significant activation of ferroptosis throughout the developmental process, offers a valuable opportunity to investigate the specific mechanisms underlying the role of ferroptosis in organ development. Our study detected the expression of Gpx4 and the accumulation of iron both in mouse mandibular incisor and developing first molar. Although Gpx4 expressed ubiquitously in both tooth germ and incisor of mouse, its abundance differs in different cell types. Results revealed an increasing iron accumulation both in odontoblast and ameloblast according to the developmental process. The spatiotemporal expression of Gpx4 is also positively linked to odontogenesis and amelogenesis. This phenomenon indicated growing ferroptotic stress (accumulating iron) and strengthening anti-ferroptosis mechanism (increasing expression of Gpx4) physiologically coexisted and may maintain a critical balance along with the differentiation and maturation of odontoblast and ameloblast. However, the discovery of changes in iron and lipid metabolism during tooth morphogenesis is not novel. In the 1930s, pioneer scientists in dental biology had already identified the presence of iron in the tooth of different animals [29–31], then found some defects of enamel in mouse is related to abnormal iron metabolism [32]. Lipid metabolism and lipid peroxidation, the other core risk factors of ferroptosis, were also described in the early stage of dental biology research [20,33,34]. Even when the risk factors of ferroptosis had been reported to participate in tooth development, there is still no reports about the exact ferroptosis related tooth developmental defects. Our results provided a new perspective to reconsider the underlying function of all these ancient studies in a comprehensive manner. They illustrated the importance of the Gpx4-dependent anti-ferroptosis pathway in managing all these already existing ferroptotic stress during tooth morphogenesis. Future *in vivo* studies utilizing transgenic mice are needed to systematically analyze the role of ferroptosis/ferroptotic stress during tooth and other organ development.

To further investigate the meaning of this precarious balance between ferroptotic stress and expression of Gpx4, we use erastin to inhibit the internalization of GSH, which is the critical substrate for Gpx4 to protect cells from lipid peroxidation. The developmental role of Gpx4 had been studied even before the ferroptosis was formally described (before 2012). In situ hybridization indicated expression of Gpx4 in all developing germ layers during gastrulation and in the somite stage in the developing central nervous system and in the heart [35], which made Gpx4 (-/-) mice die embryonically in utero by midgestation (E7.5) and are associated with a lack of normal structural compartmentalization [36]. Specific deletion of Gpx4 during developmental process were found to participate in the maturation and survival of cerebral and photoreceptor cell [37,38]. Recent years, more ferroptosis related function of Gpx4 were discovered in neutrophil [39] and chondrocyte [40] of adult mice, in which specific deletion will lead to ferroptosis-induced organ dysregulation and degeneration. Thus, it is essential to assess the biological function Gpx4-dependent ferroptotic suppressing system. In our study, erastin significantly impaired tooth morphogenesis in a dose-dependent manner within the *ex vivo* tooth germ culture model. The *ex vivo* organ culture of tooth germ is a well-established classical model for the study of tooth morphogenesis. Compared to *in vivo* study, the *ex vivo* culture of tooth germ is convenient for manipulating culture conditions and investigate factors affecting tooth germ in a high throughput way [41]. Although lacking of circulation and immune system, the *ex vivo* culture of the tooth germ, other than the traditional 2D culture of dental cells *in vitro*, can retain most properties of tooth development, like interactions among oral epithelia, mesenchyme, and stromal cells. We successfully established this model and the tooth germs from D0 to D7 are well developed (Fig S1B). To induce ferroptosis, erastin is the most used agent inhibiting GSH transport but is not stable *in vivo,* which makes *ex vivo* organ culture of tooth germ the ideal way to study the possible function of ferroptosis/ferroptotic stress in tooth morphogenesis. Moreover, according to our results, erastin treatment will not induce overactivation of apoptosis in tooth germ (Fig 5D).

The histologic analysis by HE staining showed an increased number of NLCs located in dental mesenchyme of erastin-treated tooth germ. 3D reconstruction of all the slides convinced that necrotic mainly occurred in the region of the odontoblast layer. Unlike CL-CASP3 to apoptosis, membrane localization of GSDM family proteins and MLKL to pyroptosis and necroptosis respectively, ferroptosis is a special type of cell death which have unique inducer but no special proteins reflecting its activation. Thus, ferroptosis in erastin-treated tooth germ is determined by the accumulation of iron, upregulation of 4-HNE, shrunken and condensed mitochondria, dramatically upregulated expression of *Psg2* (risk marker to ferroptosis), and mildly increased expression of *Gpx4, Slc7a11* (protective factor to ferroptosis). These results indicated that erastin could lead to abnormal tooth morphology via activating ferroptosis. Moreover, characterized by more apparent NLCs, stronger iron and 4-HNE fluorescence signal, and severer mitochondria degeneration, dental mesenchyme cell/odontoblast seems more sensitive to erastin-induced ferroptosis than that of dental epithelium cell/ameloblast.

According to our results, ferroptotic stress is physiologically increased during tooth development. Thus, rather than “initially activate ferroptosis” in tooth morphogenesis, erastin, a system x_c_^-^ inhibitor, inhibits GSH production, and induced ferroptosis in tooth germ is more likely lead by “overburdened ferroptotic stress”. As a multi-step process of cell death, the different inhibitor has the different target to suppress ferroptosis. Here in our study, regular ferroptosis inhibitors were applied in the rescue assay of impaired tooth morphogenesis. Tooth germ was treated by Fer-1, Lip-1 (radical trapping agents which inhibit the propagation of lipid peroxidation), and DFO (an iron chelator). Surprisingly, although all these inhibitors could reverse erastin-induced impairment of tooth morphogenesis, Fer-1 reduced the number of NLCs much more efficiently than lip-1 and DFO. Different from lip-1 or DFO, which only targets lipid peroxidation or labile iron accumulation, Fer-1 could both inhibit lipid peroxidation and reduce the labile iron pool in cells, notably is not consumed while inhibiting iron-dependent lipid peroxidation [42]. Sustaining double effects of Fer-1 in inhibiting ferroptosis possibly enables it effectively rescue the impairment of tooth germ than other agents.

In conclusion, by using *ex vivo* culture model of tooth germ, we identified continuing accumulation of ferroptotic stress in both odontogenesis and amelogenesis during tooth development. Activation of ferroptosis impaired tooth morphogenesis, especially within dental mesenchyme, and could be partially rescued by ferroptotic inhibitor. This study provides a promising model to effectively investigate the developmental role of ferroptosis and will broaden our knowledge about possible involvement of non-apoptotic RCD during organogenesis.

## Materials and methods

### Animals and organ culture

ICR mice were purchased from Chengdu Dossy Experimental Animals Co., Ltd (Sichuan, China). All animal work was done according to the National Institutes of Health guidebook and approved by the Committee on the Ethics of Animal Experiments of Sichuan University (WCHSIRB-D-2021-12544). The presence of a vaginal plug was used as an indication of embryonic day 0 (E0). The first mandibular molar tooth germs of E15.5 were dissected under stereomicroscope (Carl Zeiss, Germany). All the dissection steps were performed on ice. The dissected tooth germs were placed in the upper chamber of the Trans-well (3450, Corning, USA) and four tooth germs were placed on one dish. The culture medium and drugs were mixed and placed in the lower chamber. Tooth germs were cultured in DMEM/F12 supplemented with 10% fetal bovine serum, 50 U/ml penicillin/streptomycin, and 100 μg/ml ascorbic acid and incubated at 37°C and 5% CO_2_. Medium was changed every other day. Tooth germs were cultured for 5 days in the presence of different concentration of Erastin (0μM, 1.5μM, 5μM, 10μM, 20μM). Culture medium added with Fer-1(Ferrostatin-1 1μM), Lip-1(Liproxstatin-1, 200nM), DFO (Deferasirox, 100μM) were used to pretreat molar germ for 4 hours. All these small molecules were purchased from Selleck. After 5 days of cultivation, the tooth germs were fixed in 4% PFA, embedded in paraffin. Tooth germs were photographed at 0-day, 1 day, 3 day, 5 day and each tooth size (width, height and area) were measured (n=9 for each group) by using the modified method reported in the literature previously. Each experiment was repeated 3 times [43].

### Immunohistochemistry

Sections were dewaxed and rehydrated before antigen repair, and then cooled to room temperature. Sections were incubated overnight at 4°C with Gpx4 (1:200, Invitrogen, PA5-102521) in 2% bovine serum albumin (BSA)/phosphate buffered saline (PBS), pH 7.4. Negative control sections were incubated with 2% BSA/PBS. After washed, sections were then incubated with SignalStain® Boost IHC Detection Reagent (8114S, Cell Signal Technology, USA) and then washed and incubated with diaminobenzidine (DAB) to detect any reactions and were examined by light microscopy after counterstaining with hematoxylin.

### Immunofluorescence and iron probe staining

Tissue sections were treated with phosphate-buffered saline (PBS) with 0.5% Triton X-100 for permeabilization. After washing and blocking, sections were incubated with 4-HNE (1:200, Abcam, ab48506), cleaved Caspase 3 (1:200, Cell Signaling Technology, 9664) in blocking buffer (PBS and 2% bovine serum albumin) overnight at 4[. After washed, second antibody Alexa Fluor® 488 (1:200, Abcam, ab150077), Alexa Fluor® 594 (1:200, Abcam, ab150116) mixed with 4′,6-diamidino-2-phenylindole (DAPI) was applied. Incubate at room temperature for one to two hours and then wash and seal the slices.

To accurately detect the changes of iron accumulation within differently cultured tooth germs, we use an aggregation-induced emission featured iron (III) probe from ortho-substituted pyridinyl-functionalized tetraphenylethylene (TPE-o-Py) [22], which is kindly gifted by Prof. Youhong Tang. This probe displays high sensitivity and selectivity toward iron (III) detection. The recognition arises from the position isomer of ortho-substitution, and the fact that TPE-o-Py has a low acid dissociation constant (pKa) that is close to that of hydrolyzed Fe^3+^. The iron probe staining was performed as described. Briefly, TPE-o-Py was dissolved in THF (Tetrahydrofuran) before being added to PBS and diluted to working concentration of 20μM. The working solution was then placed on sections, and incubated at room temperature for half an hour and then aspirated and sealed, then a laser confocal microscope (Olympus FV3000, Japan) was used to detect the fluorescence signal. Since the TPE-o-Py probe pronounced red-shift in fluorescence emission which is positively related to the concentration of iron, low concentration of Fe^3+^ was detected under fluorescence channel of Alexa Fluor 405 (Excitation wave length: 402nm, Emission wave length: 421nm), while Alexa Fluor 594 (Excitation wave length: 590nm, Emission wave length: 617nm) for high concentration.

### Tissue preparation for transmission electron microscope

Prefixed with a 3% glutaraldehyde, then the tissue was postfixed in 1% osmium tetroxide, dehydrated in series acetone, infiltrated in Epox 812 for a longer, and embedded. The semithin sections were stained with methylene blue and Ultrathin sections were cut with diamond knife, stained with uranyl acetate and lead citrate. Sections were examined with JEM-1400-FLASH Transmission Electron Microscope.

### RNA extraction and qPCR

Total RNA of tooth germs (E15.5 cultured for 5 days) were extracted by using TRIzol™ Reagent (Invitrogen, USA) according to instruction of manufacturer. The amount and the integrity of RNA were assessed by measurement of absorbance at 260 and 280 nm. First-strand cDNA synthesis was performed with a HiScript III Q RT SuperMix for qPCR (Vazyme, China). The levels of *Gpx4, Ptgs2, Slc7a11* were measured by quantitive real-time PCR (Bio-Rad, USA) with ChamQ Universal SYBR qPCR Master Mix (Vazyme, China) and normalized to the level ofβ-Actin mRNA. These experiments were performed in triplicate. Primer sequences used in qPCR are listed below.

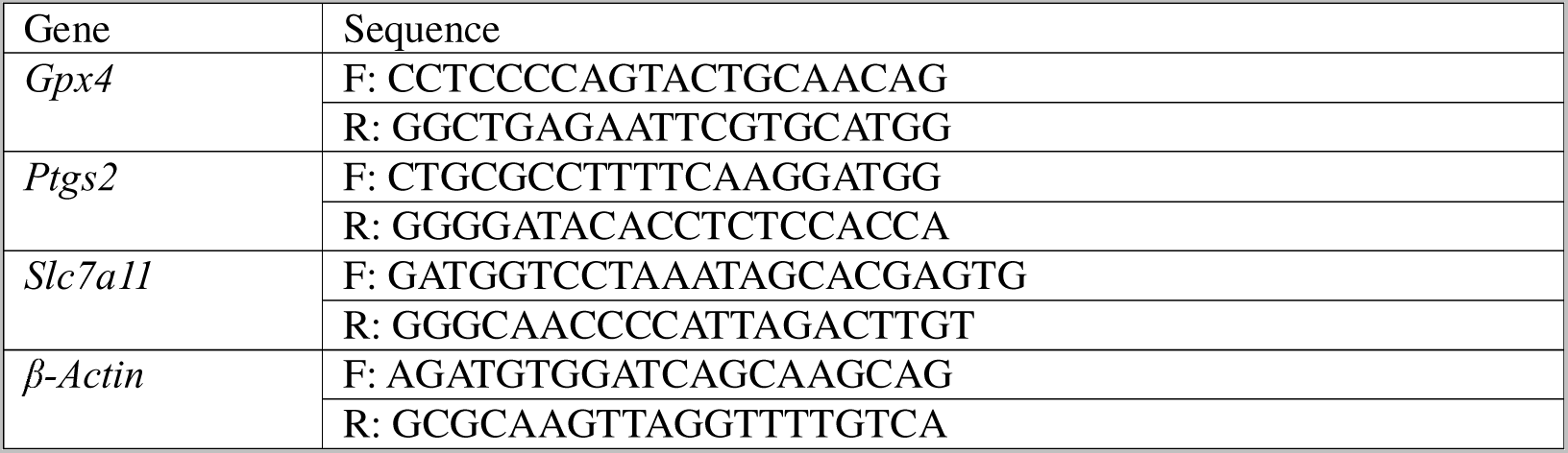

### 3D reconstruction of tooth germ sections

Sequential section and subsequent HE staining of each tooth germ were performed as described before. Digital pathological system (Olympus vs200) was used to scan all the stained sections and reconstructed by following previously described protocol [24].

### TUNEL assay

We used a One Step TUNEL Apoptosis Assay Kit (Beyotime, C1090) to detect the possible DNA damage in apoptotic cell of tooth germs. The TUNEL assay was performed following instructions.

### Statistical analysis

Analysis of Gpx4 relative expression levels were carried out by Image-pro plus7.0 (Media Cybernetics, USA). Statistical analyses were carried out using the GraphPad Prism version 8.00 (GraphPad Software, San Diego, CA, USA). All statics were shown as the arithmetic mean ± the standard error of the mean. The significance of differences between groups was tested by using the one-sample t-test. Differences were considered significant when *p<0.05*.

## Acknowledgment

This study was funded by grants from National Natural Science Foundation of China U21A20368 (L. Y.), 82101000 (H. W.), and 82201045 (F. Y.). The iron probe, TPE-*o*-Py, is kindly gifted by Prof. Youhong Tang (Flinders University, Australia) and Prof. Benzhong Tang (The Hong Kong University of Science and Technology, China).

## Author contributions

Haisheng Wang: Contributed to conception and design, acquisition, analysis, and interpretation, drafted manuscript; Xiaofeng Wang: Contributed to conception and design, acquisition, analysis, and interpretation, drafted manuscript; Liuyan Huang: Contributed to data acquisition; Chenglin Wang: Contributed to conception and design critically revised manuscript; Fanyuan Yu: Contributed to conception and design, critically revised manuscript; Ling Ye: Contributed to conception and design, critically revised manuscript. All authors gave their final approval and agree to be accountable for all aspects of the work. The authors declare no potential conflicts of interest with respect to the authorship and/or publication of this article.

---

**Figure supplement 1.**
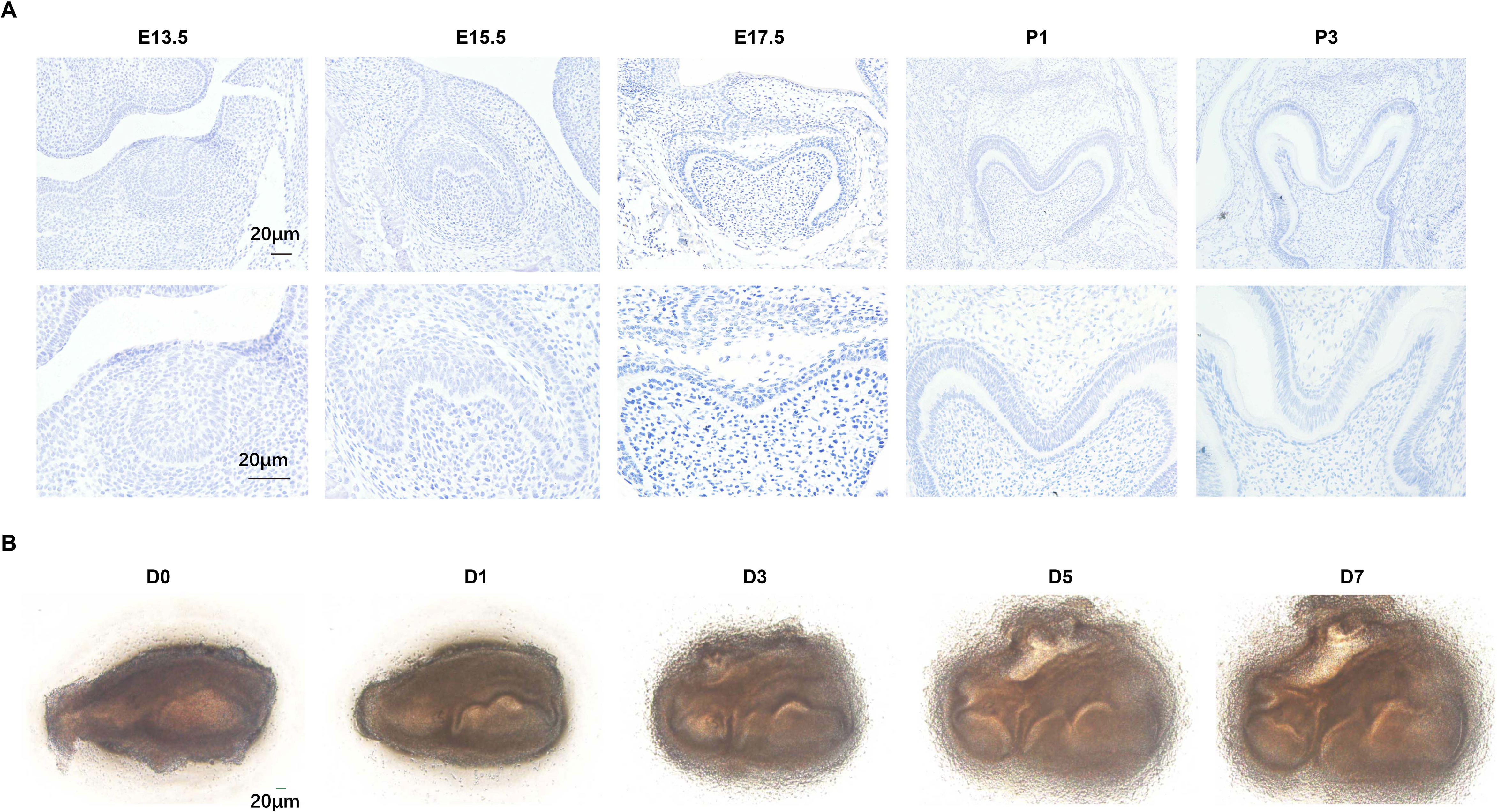
(A) Negative control of different developmental stage of molar germ in Gpx4 IHC staining. (B) *Ex vivo* culture of tooth molar germ from D0 to D7. Scar bars=20μm.

**Figure supplement 2.**
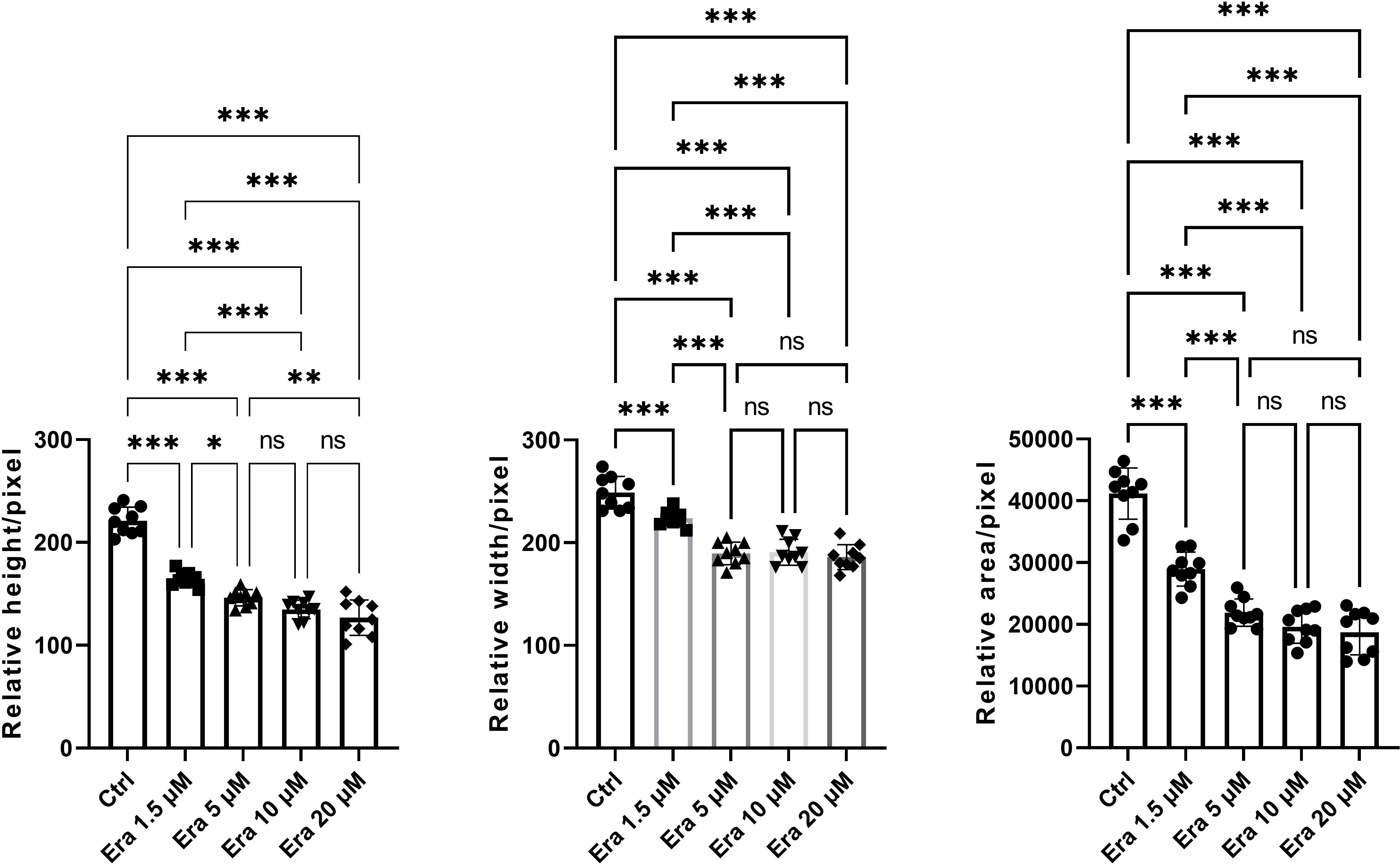
Original bar graphs of height, width, and area of erastin-treated tooth germ. *\*P<0.05, **P<0.01***P<0.001*

**Figure supplement 3.**
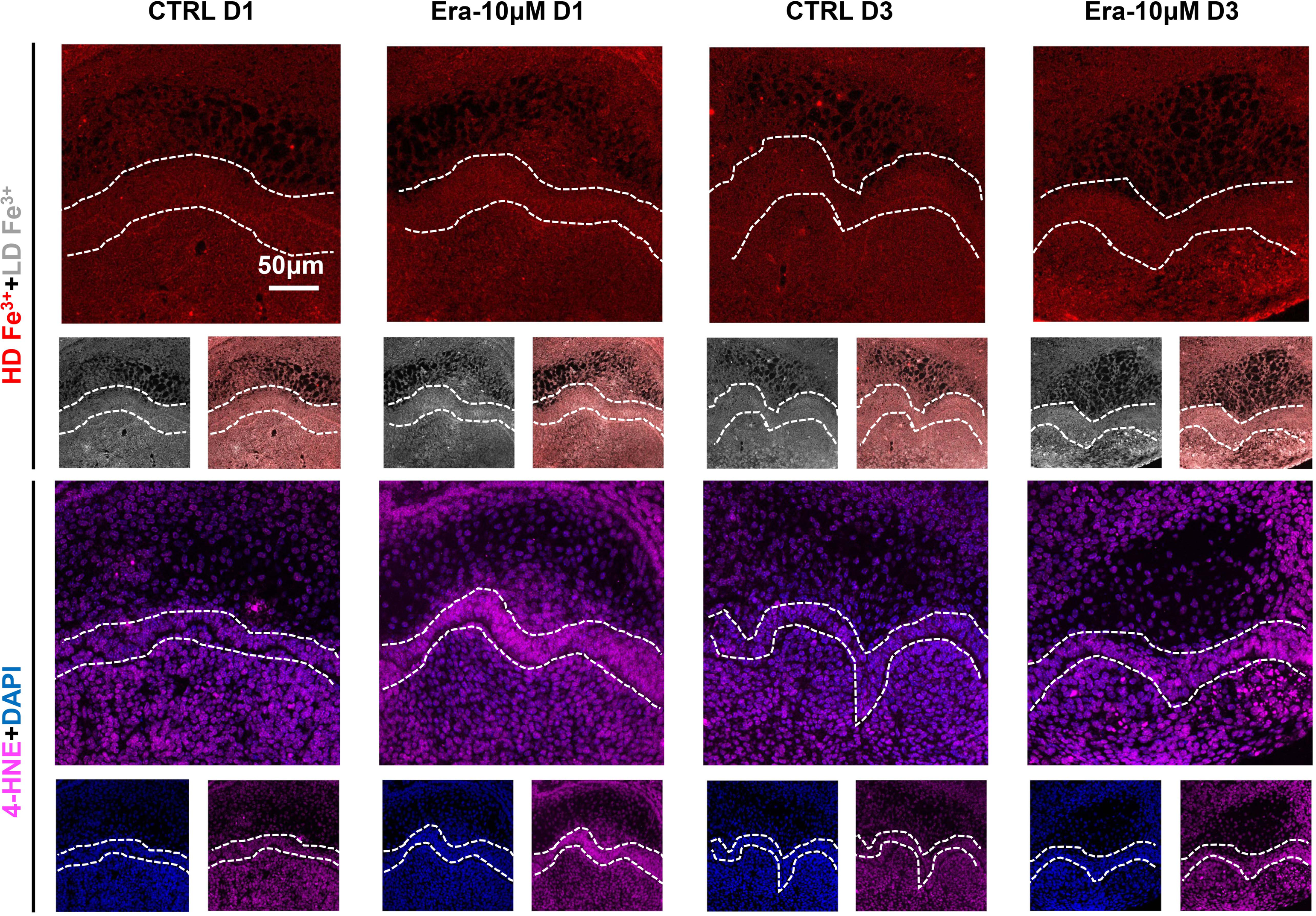
Results for iron accumulation and 4-HNE expression in Era-10μM of D1 and D3. Scar bars=50μm

**Figure supplement 4.**
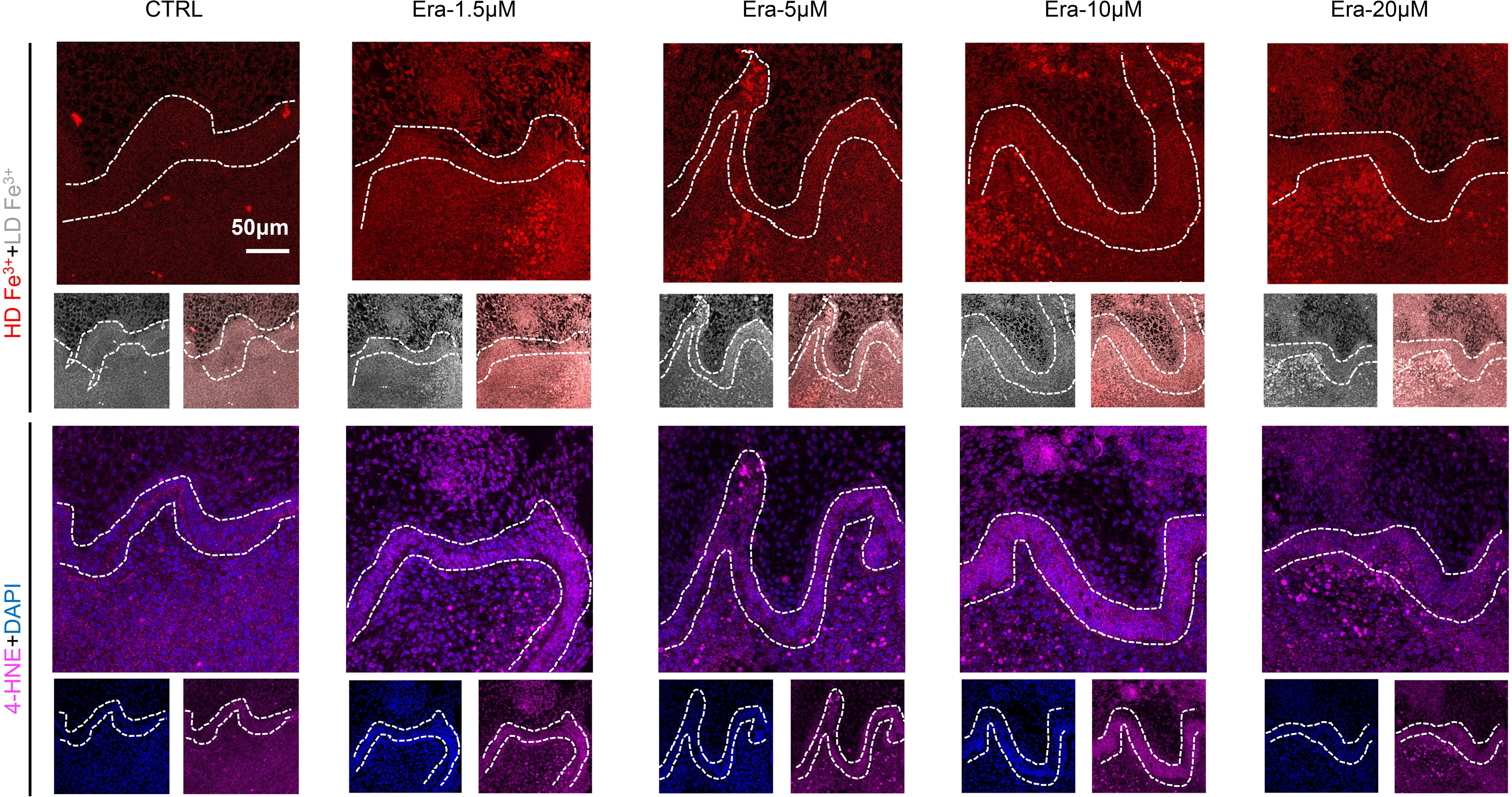
Results for iron accumulation and 4-HNE expression in Era-1.5μM, Era-5μM, Era-10μM, and Era-20μM of D5. Scar bars=50μm

**Figure supplement 5.**
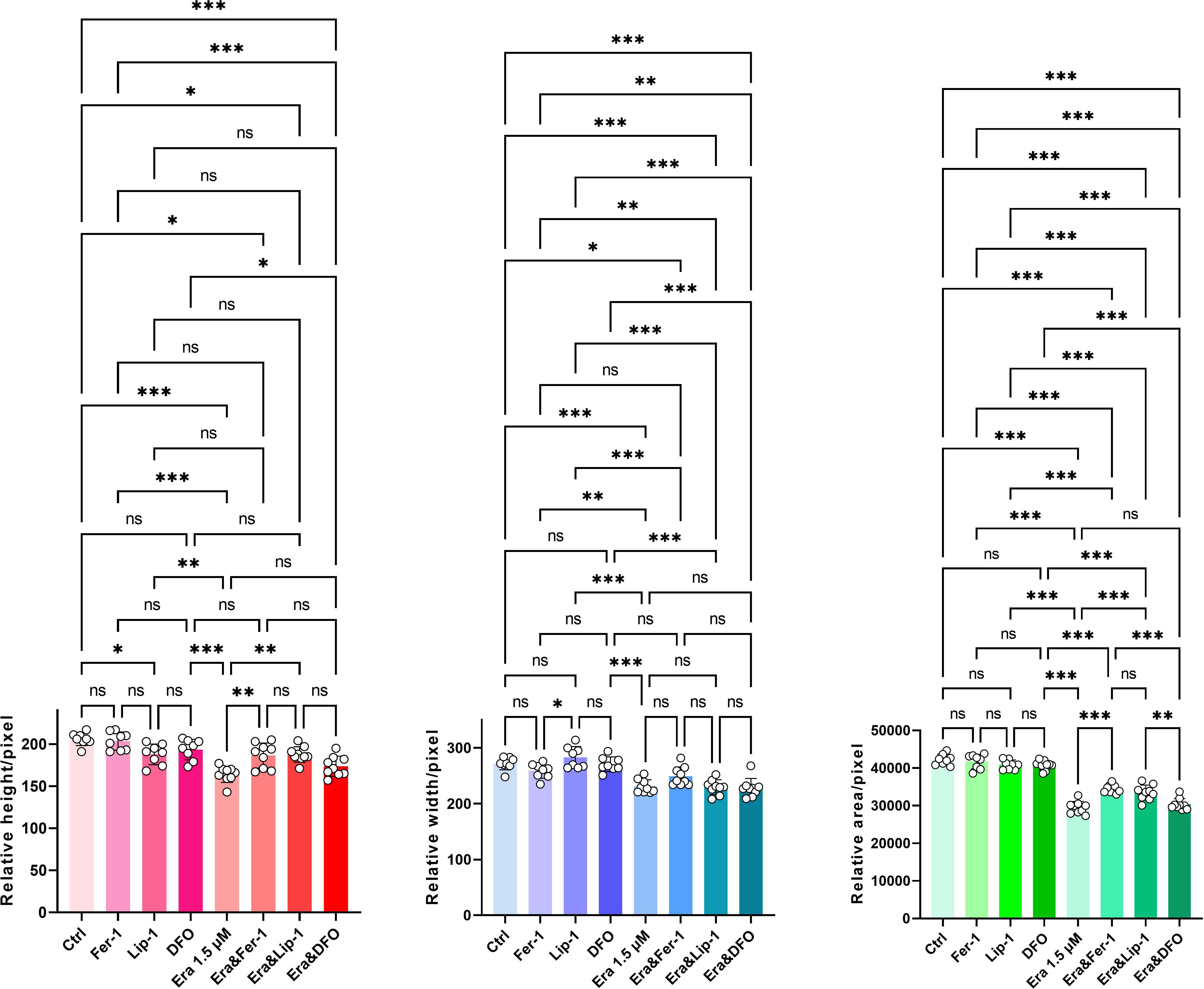
Original bar graphs of height, width, and area of tooth germ in rescue assay. *\*P<0.05, **P<0.01***P<0.001*

**Figure supplement 6.**
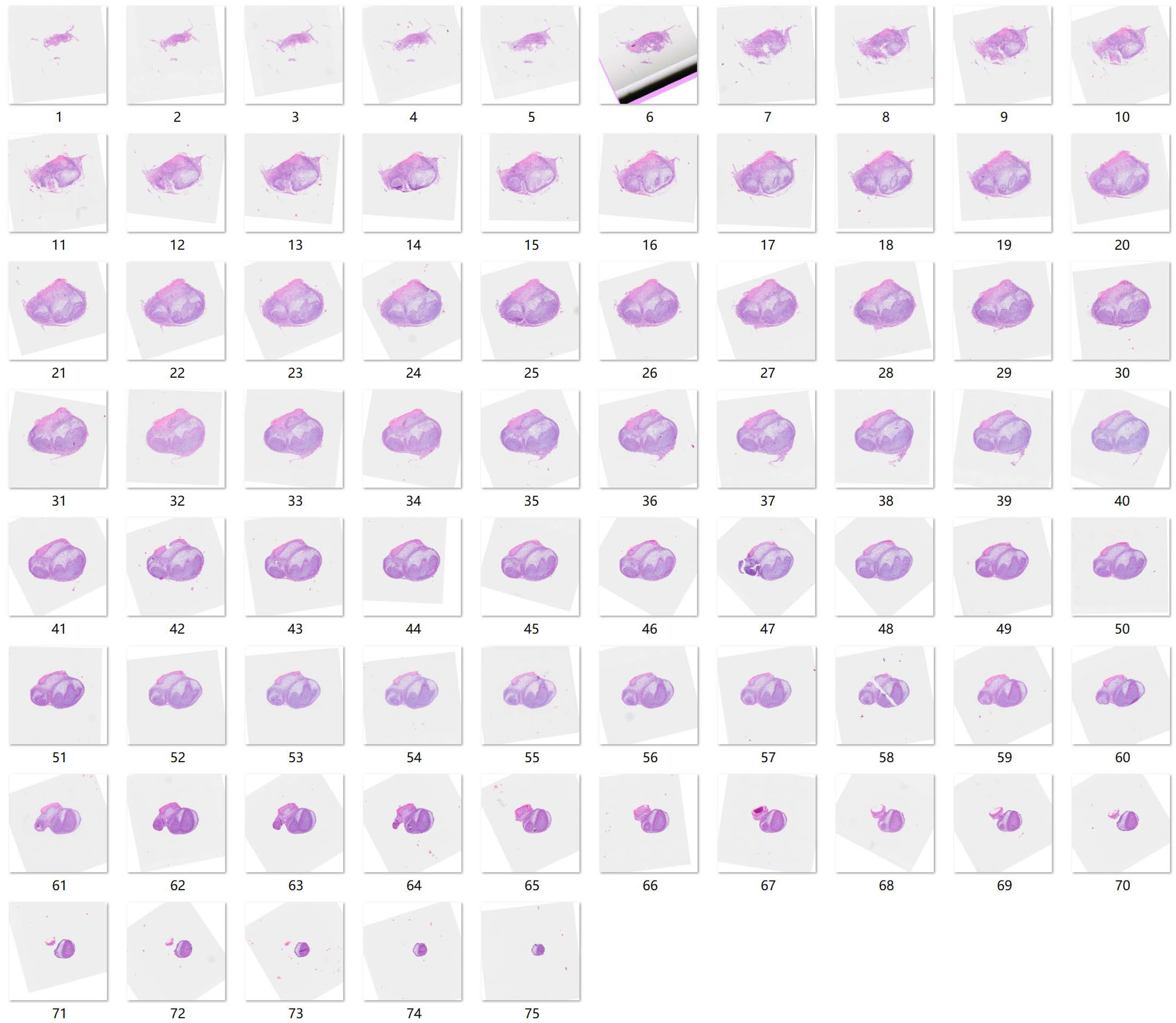
Source sequential HE slides for 3D reconstruction of CTRL

**Figure supplement 7.**
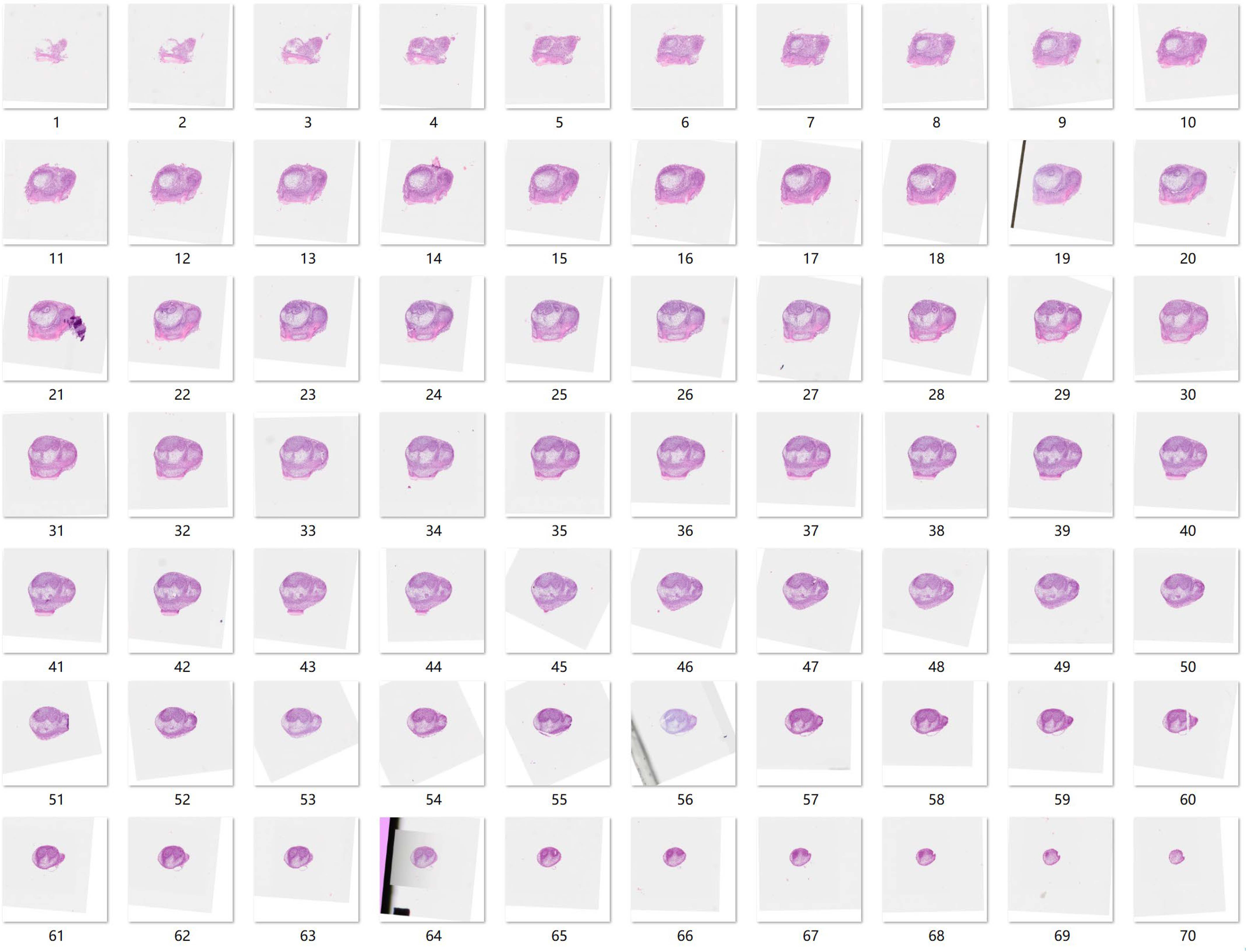
Source sequential HE slides for 3D reconstruction of Era-1.5μM

**Figure supplement 8.**
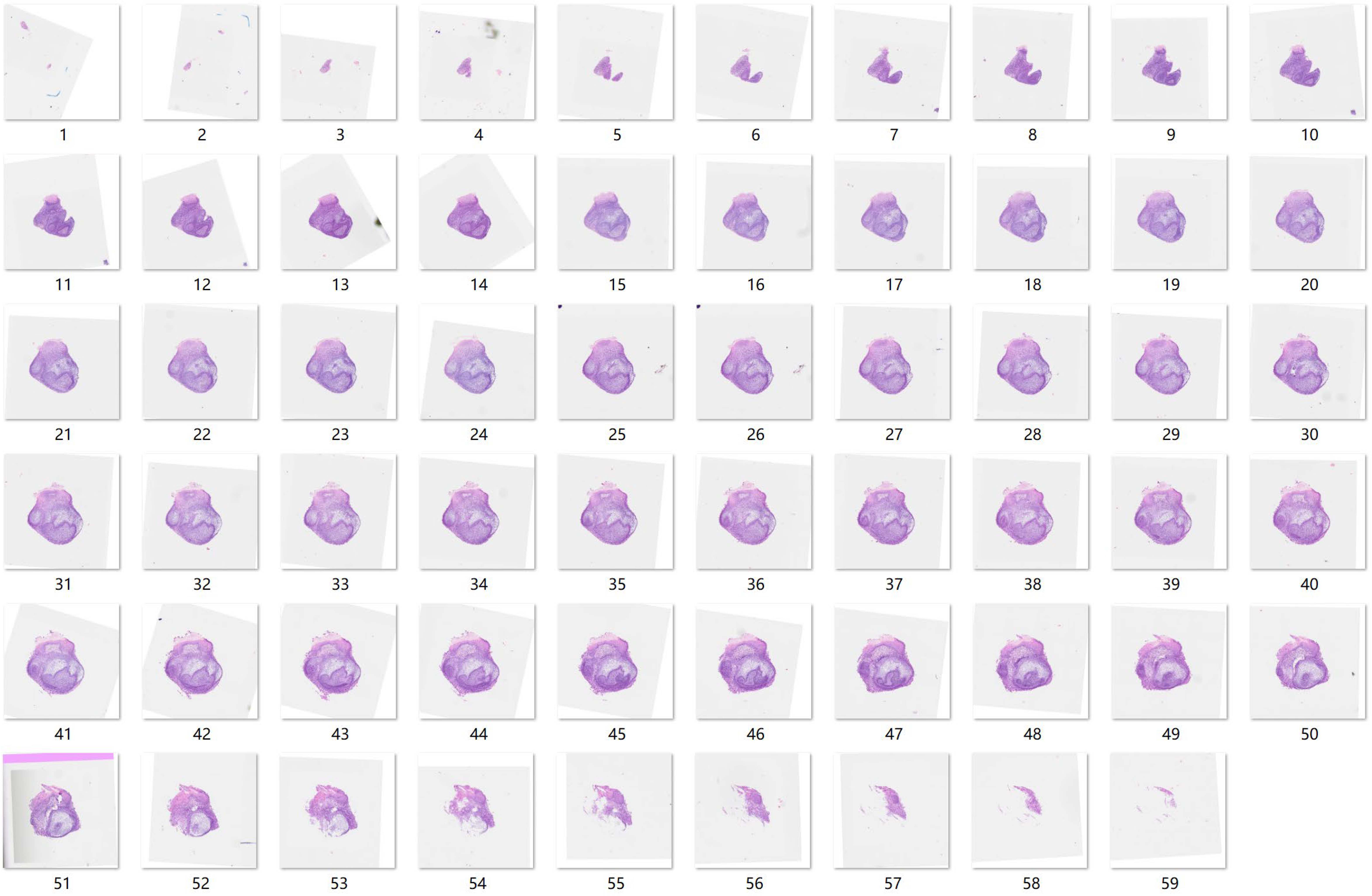
Source sequential HE slides for 3D reconstruction of Era-1.5μM+Fer

**Figure supplement 9.**
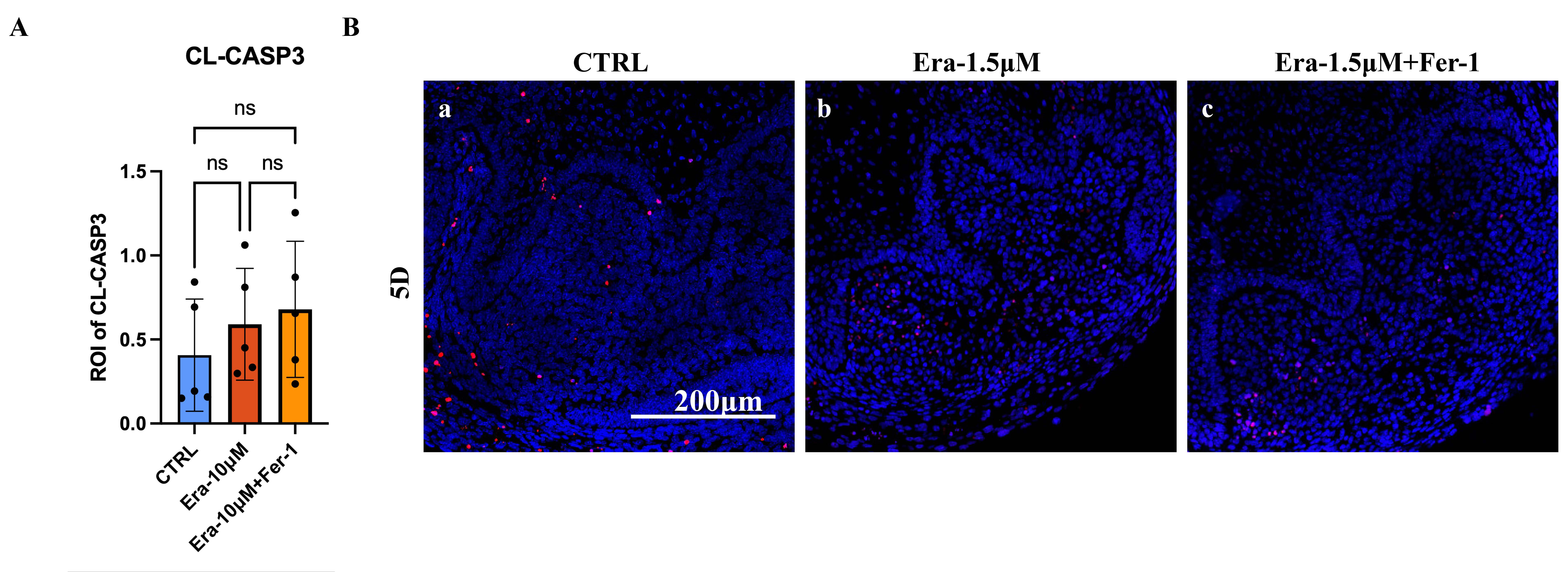
(A) Mean influence intensity of ROI of CL-CASP3 in CTRL, Era-1.5μM, and Era-1.5μM+Fer-1. Both Era-1.5μM and Era-1.5μM+Fer-1 showed slightly increased CL-CASP3 activation than that in CTRL but showed no statistical differences. (B) Results of TUNEL assay in each experimental group. DNA damage detected by TUNEL assay is similar to that of CL-CASP3, which indicated that apoptosis is not significantly activated in erastin treated tooth germ. Scar bars=200μm

## Notes

### Competing Interest Statement

The authors have declared no competing interest.

### Summary of Updates

In the current version of the manuscript, the main modifications are listed below. We replaced the data on iron concentration in the incisor with that of the developmental tooth germ (see Figure 1B). Dr. Liuyan Huang was added to the author list for her contributions during revising. She helped to address the results of the innate iron accumulation during tooth germ development. We made our abstract more concise. We modified our presentation in the manuscript. We addressed all the grammar and typo errors.

